# OmpA controls intracellular survival of *Acinetobacter baumannii* through TFEB activation and lysosomal remodeling

**DOI:** 10.64898/2026.04.18.719357

**Authors:** Irene Molina Panadero, Angela Rey Hidalgo, María José López Carballo, Celia Atalaya Rey, Manuel J Muñoz, Younes Smani

## Abstract

*Acinetobacter baumannii* is a high-priority multidrug-resistant pathogen that survives within host cells by hijacking intracellular defense pathways. Here, we identify a previously unrecognized signaling axis linking bacterial invasion to host lysosomal regulation. We show that *A. baumannii* activates calcium-independent phospholipase A2 (iPLA_2_), leading to increased lysophosphatidylcholine (LPC) production and calcium influx through the ORAI1 channel, which together drive activation and nuclear translocation of the lysosomal transcription factor EB (TFEB). Pharmacological inhibition or genetic silencing of iPLA_2_ or ORAI1 markedly impaired TFEB activation and lysosomal biogenesis. Mechanistically, we demonstrate that this pathway is initiated by the outer membrane protein A (OmpA), which promotes bacterial invasion and enhances iPLA_2_ activity, LPC production, and downstream TFEB signaling. Despite induction of lysosomal biogenesis, *A. baumannii* persists intracellularly by producing ammonia and alkalinizing the lysosomal environment, thereby counteracting host antibacterial activity*. In vivo*, infection induces activation of HLH-30, the TFEB ortholog, in *Caenorhabditis* elegans in an OmpA-dependent manner. Together, our finding define an OmpA-iPLA_2_-LPC-ORAI1-TFEB signaling axis that coordinates host lipid and calcium signaling with lysosomal responses, while revealing a bacterial counterstrategy that promotes intracellular survival.

## INTRODUCTION

The virulence of *Acinetobacter baumannii*, one of the highest-priority pathogens for the development of new antibiotics [Sati et al., 2025], relies heavily on outer membrane proteins (OMPs) that mediate adhesion, invasion, and persistence within host epithelial cells [Vila Farrés et al., 2017; Smani et al., 2013; Smani et al., *JBC* 2012; Smani et al., *PLoS One* 2012]. Adhesion represents the first and essential step of infection, followed by bacterial invasion through microfilament- and clathrin-dependent mechanisms [Smani et al., *JBC* 2012; Choi et al., 2008]. Once internalized, *A. baumannii* induces the expression of transcription factor EB (TFEB) in host cells [Parra Millán et al., 2018], a master regulator of lysosomal biogenesis and autophagy [Settembre et al., 2011].

TFEB activation enables *A. baumannii* to persist within membrane-bound vacuoles, resist intracellular clearance, and induce host cell death *in vitro* [Parra Millán et al., 2018]. To address the role of TFEB in the context of infection in a whole organism, it was further demonstrated that HLH-30, the *Caenorhabditis elegans* orthologue of TFEB, is also required for the host to cope with *A. baumannii* infection [Parra Millán et al., 2018].

The discovery that TFEB nuclear translocation is controlled by calcium through activation of the phosphatase calcineurin [Medina et al., 2015] suggested that calcium channels, particularly store-operated channels (SOCs), as well as their activators, may regulate TFEB activation via a cascade of intracellular mediators. TFEB activity is tightly regulated by phosphorylation, which retains TFEB in an inactive state in the cytoplasm; upon dephosphorylation by calcineurin, TFEB translocates to the nucleus and activates the transcription of its target genes [Medina et al., 2015].

Interestingly, *A. baumannii* has been shown to induce an increase in intracellular calcium levels through calcium release from major intracellular reservoirs, such as the endoplasmic reticulum and mitochondria, as well as by promoting extracellular calcium influx [Smani et al., 2011; Smani et al., *JBC* 2012]. Activation of SOCs via calciun-independent phospholipase A2 (iPLA₂) has revealed the involvement of lysophosphatidylcholine (LPC) in this process [Smani et al., *Nature Cell Biology* 2004]. Notably, *A. baumannii* has also been shown to increase LPC levels both *in vitro* and *in vivo* [Smani et al., *AAC* 2015].

Another aspect of TFEB activation that has been underestimated is the ability of *A. baumannii* OMPs to directly or indirectly modulate TFEB and its upstream regulators. OmpA, one of the major OMPs of *A. baumannii* [Scribano et al. 2024], is a β-barrel porin with a wide range of biologically relevant properties demonstrated both *in vitro* and *in vivo*. OmpA is involved in adherence to epithelial cells [Smith et al., 2007; Gaddy et al., 2009], translocation to the epithelial cell nucleus [Choi et al., *Cellular Microbiology* 2008], and the induction of epithelial cell death and increased mouse mortality [Choi et al., *Cellular Microbiology* 2005; Vila Farrés et al., 2017]. In humans, OmpA has also been associated with the development of pneumonia and bacteremia caused by *A. baumannii* [Sánchez-Encinales et al., 2017]. Together, these findings further support a role for OmpA in TFEB activation during *A. baumannii* infection.

In the present study, while searching for upstream regulators of TFEB activation, we identified a novel regulatory axis (iPLA₂-LPC-Ca²⁺-TFEB) that is critical for *A. baumannii* intracellular persistence. We demonstrate that this signaling pathway is initially triggered by OmpA from *A. baumannii*.

## MATERIALS AND METHODS

### Human cell culture

HeLa cells and HeLa cells stably expressing TFEB-GFP (TFEB^GFP+^cells) [Medina et al. 2015] were grown in DMEM supplemented with 10% heat-inactivated fetal bovine serum (FBS), vancomycin (50 mg/L), gentamicin (20 mg/L), amphotericin B (0.25 mg/L) (Invitrogen, Spain), and 1% HEPES in a humidified incubator at 37 °C with 5% CO_2_. Cells were routinely passaged every 3 or 4 days. Immediately before infection, cells were washed three times with pre-warmed PBS and further incubated in DMEM without FBS and antibiotics [Parra-Millán et al, 2018].

### siRNA transfection

Chemically synthesized, double-stranded small interfering RNAs (siRNAs) targeting iPLA_2_, ORAI1, and a control siRNA were purchased from Qiagen and Thermofisher, respectively. The siRNA sequences were: iPLA_2_, 5′- CTCGTTTCAACCAGAACGTTA-3’, ORAI1, 5’-CCUGUUUGCGCUCAUGAUC-3’ [Galeano-Otero et al. 2021] and control, 5′-AATTCTCCGAACGTGTCACGT-3′. HeLa cells at 40-50% confluence (1 x 10^6^ cells/well) in six-well plates, split at least 24 h prior, were transfected using HiPerfect transfection reagent (Qiagen, Spain) following a modified manufacturer’s protocol. Briefly, 6 µL of HiPerfect reagent was added to 100-µL Optimem (Invitrogen, Spain) containing 10 nM siRNA, incubated for 15 min at room temperature, and added to cells containing fresh serum-free Optimem. After 4 h at 37°C, 3 mL of DMEM with FBS and antibiotics was added. Cells were incubated for additional 48 h, and iPLA_2_ and ORAI1 expression was determined by western blotting.

### Immunofluorescence

Immunofluorescence assay was performed as described previously [Molina Panadero et al EMBO 2025]. Briefly, HeLa cells plated on coverslips were incubated with *A. baumannii* ATCC 17978 (1 x 10^7^ CFU/mL) or stauroporine (10 mg/L, apopotosis inducer) for 24 h at 37 °C, 5% CO_2_. HeLa cells were washed five times with cold PBS, fixed in methanol for 8 min at -20 °C, permeabilized with 0.5% Triton X-100, and blocked with 20% pork serum in PBS. Primary antibody, rabbit anti-iPLA_2_ (Calbiochem, USA), was applied at 1:25 in PBS containing 1% bovine serum albumin (BSA) for 2 h. After washing with PBS, coverslips were incubated with Alexa Fluor 488-conjugated goat anti-rabbit IgG (Invitrogen, Spain) at 1:50 in PBS containing 1% BSA for 1 h. Coverslips were mounted in Prolong Diamond Antifade Mounting Medium (Invitrogen, Spain) and visualized using a Leica fluorescence microscopy (Leica Microsystems, Germany).

### Western blot

HeLa cells (Control, siRNA-transfected, bromoelone lactone (BEL)-treated at 5 μM, or infected with *A. baumannii* ATCC 17978 at 1 x 10^7^ CFU/mL for 2 h) were collected, lysed in radioimmunoprecipitation assay (RIPA) buffer (Sigma, Spain) with 1 mM phenylmethylsulfonyl fluoride (PMSF) and 10% protease inhibitor cocktail (Sigma, Spain), and centrifuged at 13,000 x *g* for 20 min at 4°C. Protein concentrations were determined using the bicinchoninic acid (BCA) assay. Western blot blotting was performed as described previously [Parra Millán et al 2018]. Primary antibodies included rabbit anti-human TFEB (Cell Signaling Technology, USA), anti-human iPLA_2_ (Cayman Chemical, USA), anti-human ORAI1 (Merck, Spain), anti-human actin (Cell Signaling Technology, USA), and anti-human tubulin (Cell Signaling Technology, USA) at dilutions of 1:1,000, 1:1,000, 1:1000, 1:500 and 1:500, respectively. Horseradish peroxidase (HRP)-conjugated goat anti-rabbit IgG (Cell Signaling Technology, USA) was used as secondary antibody at dilutions of 1:5,000, 1:5,000, 1:5,000, 1:5,000 and 1:5,000, respectively). TFEB levels were normalized to actin and expressed relative to control.

### iPLA_2_ activity

iPLA_2_ activity was determined in three independent experiments as previously described [Smani et al. Nat Cell Biol 2004]. Following BEL treatment (5 and 10 μM, for 30 min) and 24 h infection with *A. baumannii* ATCC 17978 or *A. baumannii* ATCC 1798 deficient in OmpA (ΔOmpA) at 1 x 10^7^ CFU/mL, HeLa cells were collected with a cell scraper, sonicated and centrifuged at 10,000 x *g* for 15 min at 4 °C. The supernatant was assayed the same day using a modified cPLA_2_ kit (cPLA_2_ assay kit; Cayman chemical, USA). Samples were incubated with arachidonoyl Thio-PC in a calcium-free buffer (300 mM NaCl, 0.5% Triton X-100, 60% glycerol, 4 mM EGTA, 10 mM HEPES, pH 7.4, 2 mg/mL BSA) for 1 h at 20 °C. The reaction was stopped with DTNB for 5 min, and absorbance was determined at 405 nm using a standard plate reader (POLARstar Omega, BMG Labtech GmbH, Germany).

### Cyclooxygenase-2 activity and prostaglandin E2 measurement

Cyclooxygenase-2 (Cox-2) activity was determined in three independent experiments using the Cox-activity assay kit (Cayman Chemical, USA). After BEL (10 μM) or Cox inhibitor NS (50 μM) treatment for 30 min and 24 h infection with *A. baumannii* ATCC 17978 1 x 10^7^ CFU/mL, HeLa cells were collected with a cell scraper, sonicated and centrifuged at 10,000 x *g* for 15 min at 4 °C. Supernatants were assayed day following kit instructions.

Prostaglandin E2 (PGE2) levels were quantified in three independent experiments using the PGE2-EIA monoclonal kit (Cayman Chemical, USA) from culture supernatants collected after the same treatments.

### LPC production

LPC levels were determined in three independent experiments following the instructions of LPC assay kit (Azwell, Japan). After BEL treatments (10 μM, 30 min) and 24 h infection with *A. baumannii* ATCC 17978 or ΔOmpA at 1 x 10^7^ CFU/mL, HeLa cells culture supernatants were collected for analysis.

### TFEB activation

TFEB^GFP+^cells seeded in 24-well plates were infected for 2 h with *A. baumannii* ATCC 17978 or ΔOmpA at 1 x 10^7^ CFU/mL in the presence or absence of ORAI1 inhibitor GSK 7975A (20 μM, 30 min) (Aobious, Gloucester, MA, United States) [Galeano et al 2021], EGTA (4 mM, 30 min) or treated with LPC (0.5 μM, 1 h). After washing three time the cells by PBS 1X, total GFP fluorescence intensity was measured using a plate-reading fluorometer (TECAN SPARK 10M, Austria). Increase fluorescence indicated TFEB-GFP accumulation and activation.

### TFEB nuclear translocation assay

TFEB^GFP+^cells were seeded in 24-well plates and infected with *A. baumannii* ATCC 17978 at 1 x 10^7^ CFU/mL in the presence or absence of BEL (5 μM, 30 min) for 2 h at 37 °C, 5% CO_2_ in three independent experiments. HeLa cells were washed three times with pre-warmed PBS. At least five image fields per well were acquired using ZOE Fluorescent Cell Imager (BioRad, Spain). Percentage of TFEB nuclear translocation was calculated as: (number of HeLa cells with nuclear TFEB / total number of HeLa cells) × 100.

### Lysosome staining

Lysosomes were visualized in HeLa cells (1 × 10^5^ cells/well) after 2 h infection with *A. baumannii* ATCC 17978 or ΔOmpA at 1 x 10^7^ CFU/mL in the presence or absence of GSK 7975A (20 μM, 30 min). Control and infected HeLa cells were incubated with LysoTracker Red (75 nM, 90 min) and MitoTracker Green (250 nM, 45 min) a marker of mitochondria, (Invitrogen, Spain), and imaged using a ZOE Fluorescent Cell Imager (BioRad, Spain).

### Lysosomal and bacterial pH determination

Lysosomal pH was determined in three independent experiments using LysoSensor Yellow/Blue dextran staining (Thermo Fisher Scientific, Spain). Briefly, HeLa cells were infected with *A. baumannii* ATCC 17978 or ΔOmpA at 1 x 10^7^ CFU/mL and stained with 0.5 mg/mL LysoSensor Yellow/Blue dextran for 2 h and visualized by STELLARIS 5 fluorescence microscopy (Leica Microsystems CMS GmbH, Germany). Yellow and blue fluorescence signals correspond to acidic and neutral lysosomal compartments, respectively. To quantify lysosomal pH, a standard calibration curve (pH 3-7) was generated in the of nigericin and monesin (Merck, Spain), ionophores used to equilibrate intracellular with extracellular pH.

For pH determination in bacterial cultures, *A. baumannii* ATCC 17978 and ΔOmpA strains were grown in different LB media at pH 4.8 and 7.1, inoculated at 0.1 OD_600_ and incubated a 37°C for 24 h. pH measurements were taken at defined time points for culture and control media with without bacteria.

### Ammonia and fumarate production

Ammonia and fumarate production were determined in three independent experiments using the Ammonia Assay Kit (Merck, Spain) and EnzyChrom Fumarate Assay Kit (Bioassay System, USA), respectively, according to the manufacturer’s instructions. Briefly, after infection of HeLa cells with *A. baumannii* ATCC 17978 or ΔOmpA at 1 x 10^7^ CFU/mL for 2 h, cells lysates were collected to measure ammonia levels.

In addition, ammonia production by *A. baumannii* ATCC 17978 or ΔOmpA was also evaluated in three independent experiments using the same kit assay. Briefly, both strains were grown overnight in LB medium at 37° C, diluted 1/10 in DMEM and incubated for 2 h. Bacterial cultures were then centrifugated at 4000 x *g* for 3 min, washed with two time with PBS, and then resuspended in the assay buffer of the ammonia kit, and then followed the kit protocol. The bacteria were incubated for 2 h at 37 °C with shaking conditions, and ammonia levels were determined at the specified time point.

### Bacterial internalization assay

HeLa cells were infected with *A. baumannii* ATCC 17978 or ΔOmpA at 1 x 10^7^ CFU/mL for 2, 4 or 8 h in the presence or absence of bafilomycin (0.8 μM, 30 min) at 37°C and 5% CO_2_. After infection, HeLa cells were washed five times with pre-warmed PBS and incubated for an additional 30 min in DMEM containing 256 mg/L gentamicin to kill extracellular bacteria (the gentamicin MIC for ATCC 17978 and ΔOmpA is 0.5 mg/L). Cells were then washed three times with PBS to remove residual antibiotic and lysed with 0.5% Triton X-100. Serial dilutions of the lysates were plated onto LB agar and incubated at 37°C for 24 h to enumerate colonies of developed and determine the number of bacteria that had invaded HeLa cells.

### *C. elegans* strain and growth conditions

For *C. elegans* experiments, *E. coli* OP50, *A. baumannii* ATCC 17978 and *A. baumannii* ATCC 17978 ΔOmpA strains were used. All strains were grown overnight at 37°C in LB medium; the ATCC 17978 ΔOmpA strain was grown in the same conditions in the presence of ticarcillin. Stationary-phase cultures were diluted to 1 x 10^7^ CFU/mL using the McFarland standard, and 100 µL aliquots were seeded onto 35-mm nematode growth medium (NGM) plates [Stiernagle 2006]. Plates were incubated at room temperature for 24 h prior to use. The *C. elegans* strain used in this work was MAH240 {*sqIs17 [Phlh-30::hlh-30::GFP; rol-6(su1006)]*} (CGC, NIH, USA) [Lapierre et al 2013].

### C. elegans HLH-30::GFP nuclear translocation assays

HLH-30::GFP nuclear translocation was quantified in first-day adult worms grown at 20°C in NGM plates seeded with OP50. Worms were subsequently exposed to *A. baumannii* ATCC 17978 or ATCC 17978 ΔOmpA for 24, 28 and 32 h at 20°C. After exposure, worms were then incubated on ice for 15-30 minutes to induce paralysis. HLH-30::GFP subcellular localization was visualized using a Leica M205 fluorescence microscope (Leica Microsystems, Switzerland).

### Statistical analysis

Group data are presented as means ± standard errors of the means (SEM). The Student t test was used to determine differences between means using the GraphPad Prism 9 (version 9.3.1; GraphPad Software, LLC). *P* < 0.05 was considered significant.

## Results

### *A. baumannii* increases iPLA_2_ activation and its mediators

To explore whether exposure to *A. baumannii* triggers cellular stress signaling, we examined the effect of *A. baumannii* ATCC 17978 on iPLA_2_ expression and activation, as well as on its downstream mediators Cox-2 and PGE2. Fluorescence microscopy images revealed a significant increase in iPLA_2_ expression (green fluorescence) in HeLa cells infected with *A. baumannii* or treated with staurosporine (a known inducer of cell death), compared with control cells (**Figure 1A**). This increase in expression was accompanied by a twofold increase in iPLA_2_ enzymatic activation, as demonstrated by western blot and enzymatic activity assays (**Figure 1B and C**). Importantly, iPLA_2_ activity was significantly reduced in a dose-dependent manner in the presence of the irreversible iPLA_2_ inhibitor BEL (*P*=0.05 and *P*=0.042) (**Figure 1C**). Because Cox-2 activity and PGE2 production are known downstream of iPLA_2_ activation [Lüthi and Marin 2007], we assessed their levels following infection. *A. baumannii* significantly increased both Cox-2 and PGE2 levels (*P*=0.001 and *P*=0.004), and these increases were markedly attenuated by BEL and Cox-inhibitor NS (**Figure 1D and E**). Together, these data demonstrate that *A. baumannii* activates the iPLA_2_ pathway, leading to increased Cox-2 activity and PGE2 production.

**Figure 1.**
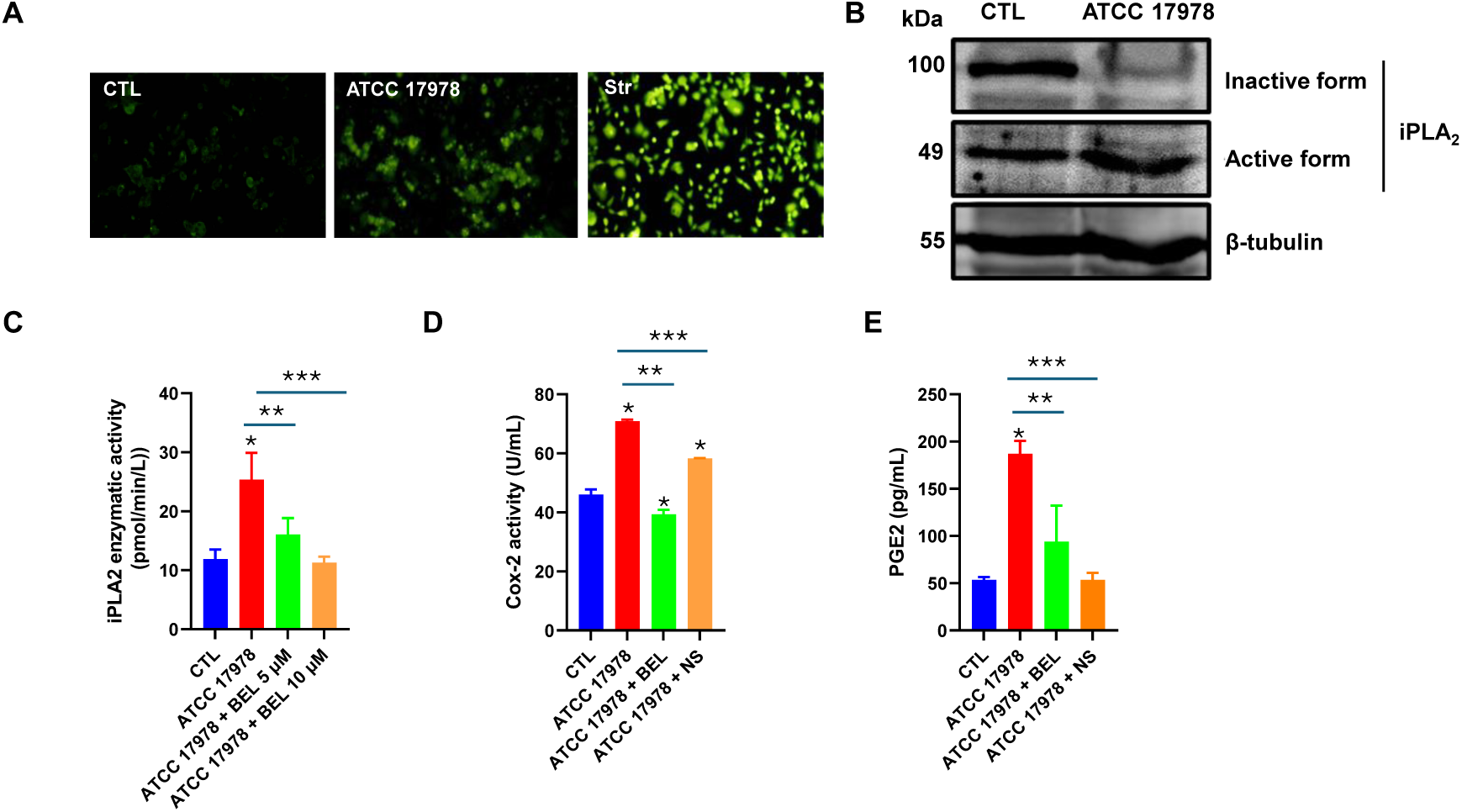
*A. baumannii* increases the expression and activity of iPLA_2_ and associated mediators. (A) Immunostaining of iPLA_2_ in HeLa cells infected with the ATCC 17978 strain for 24 h. **(B)** Immunoblot analysis of inactive and active forms of iPLA_2_ in HeLa cells infected with the ATCC 17978 strain for 24 h. **(C)** Enzymatic activity of iPLA_2_ in HeLa cells infected with the ATCC 17978 strain for 24 h, in the presence or absence of BEL. \**P*=0.025 for ATCC 17978 vs. CTL, \*\**P*=0.05 for ATCC 17978 vs. ATCC 1797 8 + BEL (5 µM), \*\*\**P*=0.042 for ATCC 17978 vs. ATCC 17978 + BEL (10 µM). **(D)** Cox-2 enzymatic activity in HeLa cells infected with the ATCC 17978 strain for 24 h, in the presence or absence of BEL and NS. (\**P*=0.0011 for ATCC 17978 vs. CTL, \**P*=0.049 for CTL vs. ATCC 17978 + BEL (5 µM), \**P*=0.0096 for CTL vs. ATCC 17978 + NS, \*\**P*=0.0005 for ATCC 17978 vs. ATCC 17978 + BEL (5 µM), \*\*\**P*=0.001 for ATCC 17978 vs. ATCC 17978 + NS. (**E**) PGE2 production in HeLa cells infected with the ATCC 17978 strain for 24 h, in the presence or absence of BEL and NS. \**P*=0.004 for ATCC 17978 vs. CTL, \*\**P*=0.0417 for ATCC 17978 vs. ATCC 17978 + BEL (5 µM), \*\*\**P*=0.0015 for ATCC 17978 vs. ATCC 17978 + NS. CTL: control; Str: staurosporine; BEL: bromoelone lactone; NS: cyclooxygenase inhibitor; Cox-2: cyclooxygenase-2; PGE2: prostaglandin 2.

### iPLA_2_ regulates TFEB activation during *A. baumannii* infection

Since TFEB is a master regulator of lysosomal biogenesis and autophagy [Settembre et al 2011], we suggest that iPLA_2_ activation observed in **figure 1** may act as an upstream signaling event during bacterial infection. To test this hypothesis, we first examined whether iPLA_2_ inhibition affects TFEB expression. Western blot analysis demonstrated that treatment with BEL significantly reduced the TFEB protein levels compared with cells infected with *A. baumannii* alone (*P*=0.041) (**Figure 2A**). Furthermore, fluorescence microscopy analysis of TFEB^GFP+^cells revealed that *A. baumannii* infection induced significant TFEB nuclear translocation compared with control cells (*P*=0.0162). Importantly, BEL treatment markedly attenuated this nuclear translocation (*P*=0.0497) (**Figure 2B**). These results indicate that iPLA_2_ activity not only regulates TFEB protein expression but also promotes its functional translocation to the nucleus. To confirm the specificity of the pharmacological inhibition results, we used siRNA-mediated iPLA_2_ knockdown. While scrambled control siRNA had no significant effect, silencing iPLA_2_ resulted in a significant reduction in TFEB protein levels (*P*<0.0012) (**Figure 2C**). This genetic approach validates that iPLA_2_ specifically modules TFEB expression during *A. baumannii* challenge.

**Figure 2.**
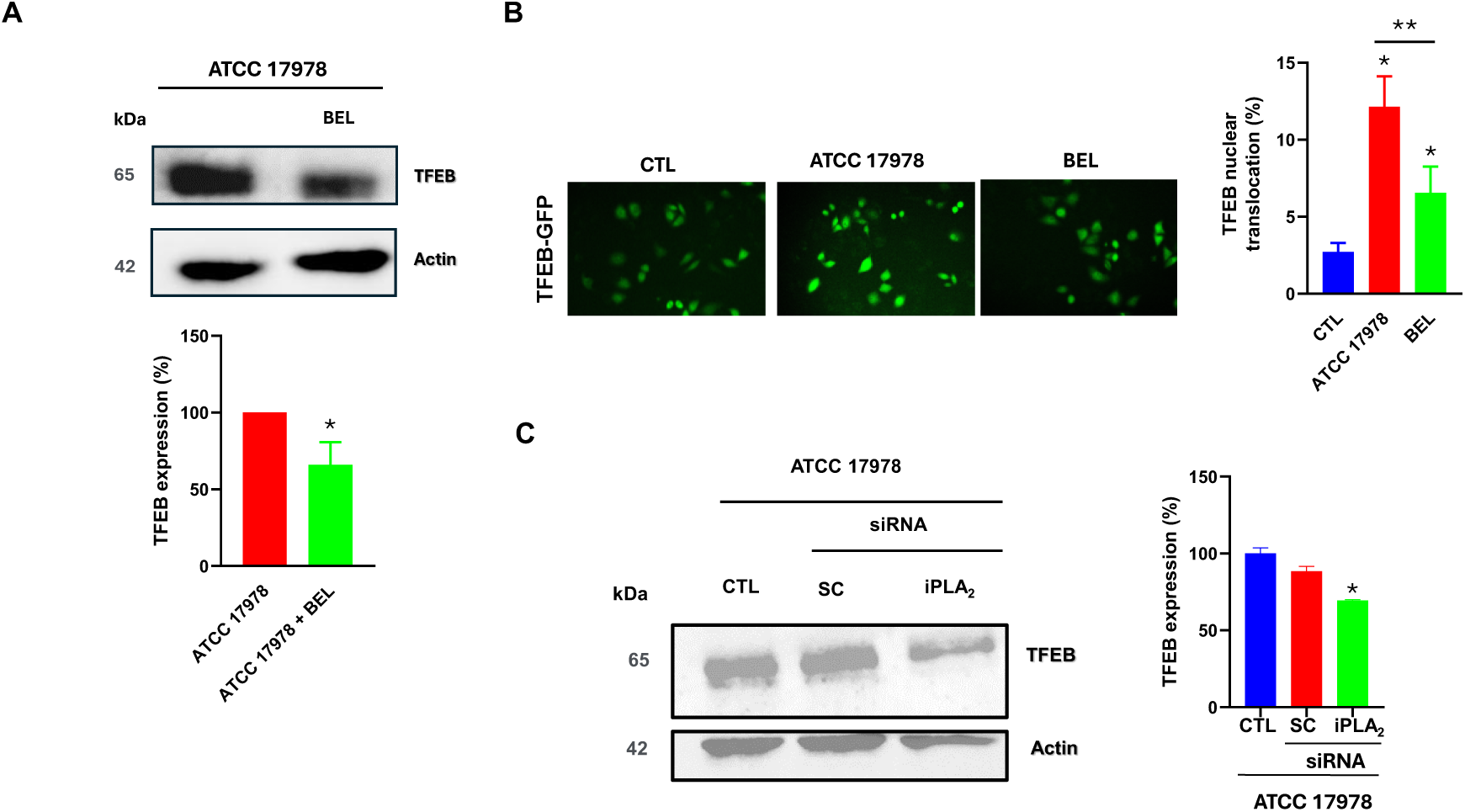
iPLA_2_ is involved in the TFEB production by *A. baumannii*. **(A)** Immnunoblot analysis of TFEB in HeLa cells infected with the ATCC 17978 strain for 2 h in the presence or absence of BEL (5 µM). \**P*=0.041 for ATCC 17978 vs. ATCC 17978 + BEL. **(B)** TFEB nuclear translocation in TFEB^GFP+^cells infected with the ATCC 17978 strain for 2 h in the presence or absence of BEL (5 µM). \**P*=0.0162 for CTL vs. ATCC 17978, \**P*=0.0452 for CTL vs. ATCC 17978, \*\**P*=0.0497 for ATCC 17978 vs. ATCC 17978 + BEL. **(C)** Immunoblot analysis of TFEB in HeLa cells transfected with scrambled or iPLA_2_ siRNA for 48 h and infected with the ATCC 17978 strain for 2 h. \**P*=0.0012 for CTL vs. siRNA of iPLA_2_. BEL: bromoelone lactone; Sc: scramble.

### iPLA_2_-derived LPC contributes to TFEB activation

LPC is generated following iPLA_2_ activation [Smani et al 2004]. To determine whether LPC links iPLA_2_ to TFEB activation during *A. baumannii* infection, we first measured LPC levels in infected cells, with or without BEL treatment. Infection with *A. baumannii* ATCC 17978 strain resulted in an approximately threefold increase in LPC concentration compared with control cells (*P*<0.0001). Treatment with BEL significantly reduced LPC levels produced by 33% during infection (*P*=0.048) (**Figure 3A**), indicating that iPLA_2_ regulates the LPC generation in this context. Next, we explored whether LPC directly activates TFEB. Cells treatment with exogenous LPC induced approximately a 25% increase in TFEB activation compared with baseline (**Figure 3B**). Together, these findings indicate that LPC acts as a bioactive lipid mediator capable of promoting TFEB activation, thereby linking iPLA_2_ signaling to lysosomal transcriptional regulation.

**Figure 3.**
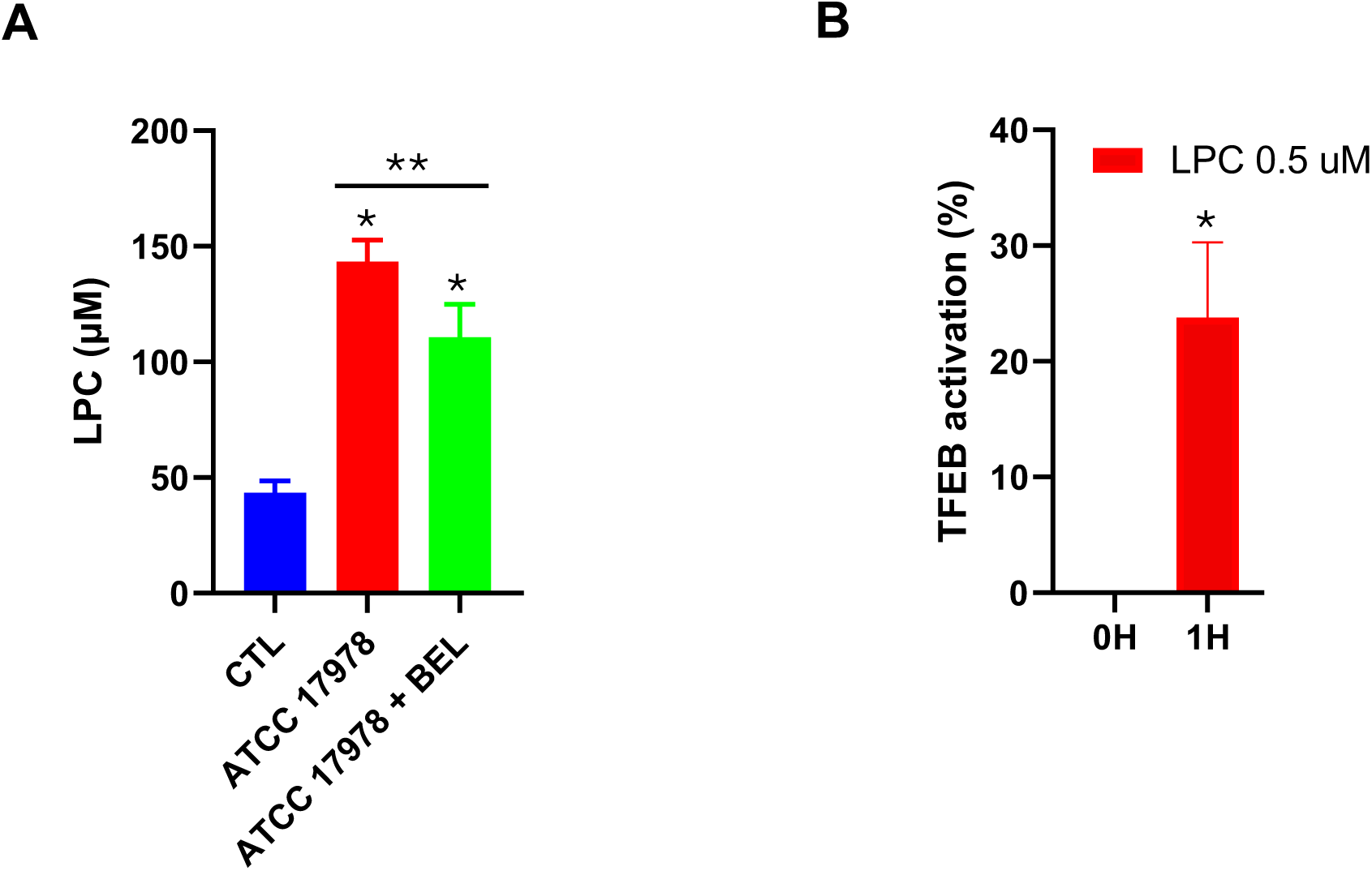
iPLA_2_-derived chemotactic factor LPC is involved in TFEB activation by *A. baumannii*. **(A)** LPC production in HeLa cells infected with the ATCC 17978 strain for 24 h, in the presence or absence of BEL. \**P*<0.0001 for ATCC 17978 vs. CTL, \**P*=0.032 for CTL vs. ATCC 17978 + BEL, \*\**P*=0.048 for ATCC 17978 vs. ATCC 17978 + BEL. **(B)** TFEB activation in TFEB^GFP+^cells treated with LPC (0.5 µM) for 1 h. \**P*<0.0001 for 0 h vs. 1 h. CTL: control; BEL: bromoelone lactone; LPC: lysophosphatidylcholine.

### Calcium influx via ORAI1 is required for TFEB activation

TFEB nuclear translocation is regulated by calcium-dependent activation of phosphatase calcineurin [Medina et al., 2015]. Based on this, we suggested that calcium influx, particularly through SOCs, may regulate TFEB activation during *A. baumannii* infection. To test this hypothesis, we first pharmacologically inhibited the SOC by ORAI1 inhibitor GSK 7975A. Treatment with the ORAI1 inhibitor significantly reduced TFEB activation in infected TFEB^GFP+^cells from approximately 28% to 20% (*P*=0.05) (**Figure 4A)**. Similarly, extracellular calcium chelation by EGTA reduced TFEB activation in infected TFEB^GFP+^cells compared with untreated infected cells (*P*<0.0001) (**Figure 4B**). To genetically validate these findings, ORAI1 expression was silenced using siRNA (**Figure S1**). ORAI1 knockdown significantly reduced TFEB protein levels compared with scrambled siRNA and control cells (*P*=0.0093) (**Figure 4C**), demonstrating that ORAI1 is a main calcium channel involved in regulating TFEB expression and activity in this model. Furthermore, infection of HeLa cells with *A. baumannii* leads to a marked increase in lysosomal labeling (red fluorescence), consistent with enhanced lysosomal biogenesis. This increase was attenuated in the presence of the ORAI1 inhibitor (**Figure 4D**). Collectively, these findings demonstrate that calcium influx through ORAI1 is an essential upstream regulator of TFEB-dependent lysosomal biogenesis during *A. baumannii* infection.

**Figure 4.**
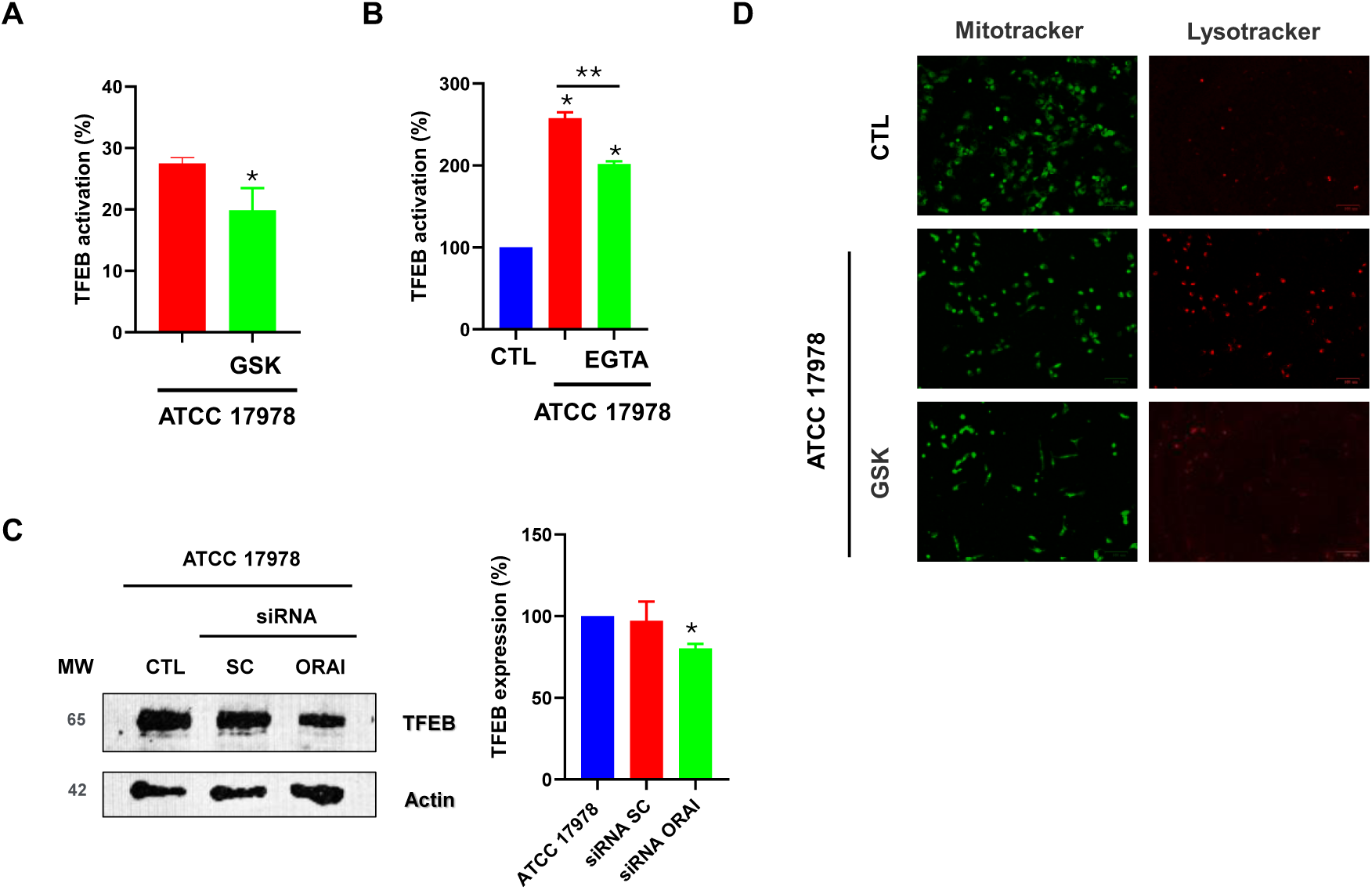
Calcium modulates TFEB activation by *A. baumannii*. **(A)** Inhibition of calcium entry through ORAI channel reduces TFEB activation in TFEB^GFP+^cells infected with the ATCC 17978 strain for 2 h. \**P*=0.05 for ATCC 17978 vs. ATCC 17978 + GSK. 7975A **(B)** Chelation of intracellular calcium with EGTA reduces TFEB activation in TFEB^GFP+^cells infected with the ATCC 17978 strain for 2 h. \**P*<0.0001 for CTL vs. ATCC 17978, \**P*<0.0001 for CTL vs ATCC 17978 + EGTA, \*\**P*<0.0001 for ATCC 17978 vs. ATCC 17978 + EGTA. **(C)** Immunoblot analysis of ORAI and TFEB in HeLa cells transfected with scrambled or iPLA_2_ siRNA for 48 h and infected with the ATCC 17978 strain for 2 h. \**P*=0.0093 for ATCC 17978 vs. siRNA ORAI. **(D)** Lysosomal analysis in HeLa cells infected with the ATCC 17978 strain for 2 h. Acidic organelles were labeled with LysoTracker Red and mitochondria were labeled with MitoTracker Green. CTL: control, EGTA: Ethylene glycol-bis(β-aminoethyl ether)-N,N,N′,N′-tetraacetic acid tetrasodium salt.

### *A. baumannii* promotes intracellular survival through ammonia production and environmental alkalinization

We next explored how *A. baumannii* survives intracellularly despite enhanced lysosomal biogenesis. Infection with *A. baumannii* resulted in a significant increase in ammonia and lysosomal pH levels compared with uninfected control cells (*P*=0.0011 and *P*=0.0164, respectively) (**Figure 5A and C**). Furthermore, treatment of infected cells with bafilomycin, an inhibitor of the vacuolar H^+^-ATPase responsible for lysosomal acidification, significantly enhanced bacterial replication. *A. baumannii* replication increased ≈ 60-fold at 8 h in the presence of bafilomycin, compared with a 20-fold increase in untreated infected cells (**Figure 5B**). These findings indicate that lysosomal acidification restricts *A. baumannii* growth and that *A. baumannii* may counteract this defense mechanism, at least in part, through the production of ammonia. Given the ability of *A. baumannii* to survive and replicate within host cells, we next examined its capacity to tolerate and modulate acidic environments. When cultured under acidic conditions (initial pH 4.8), *A. baumannii* progressively increased the pH, reaching nearly pH 9 after 24 h of growth (**Figure 5D**), although their growth dynamic is different in the first hours of bacterial growth (**Figure 5E**). In contrast, in the absence of bacteria, the pH of the culture medium (initially pH 7.1 or 4.8) remained relatively stable over the same period (**Figure S2**). These results demonstrate that *A. baumannii* possesses a strong capacity to alkalinize its environment, likely facilitating intracellular survival by survival by counteracting host-mediated acidification.

**Figure 5.**
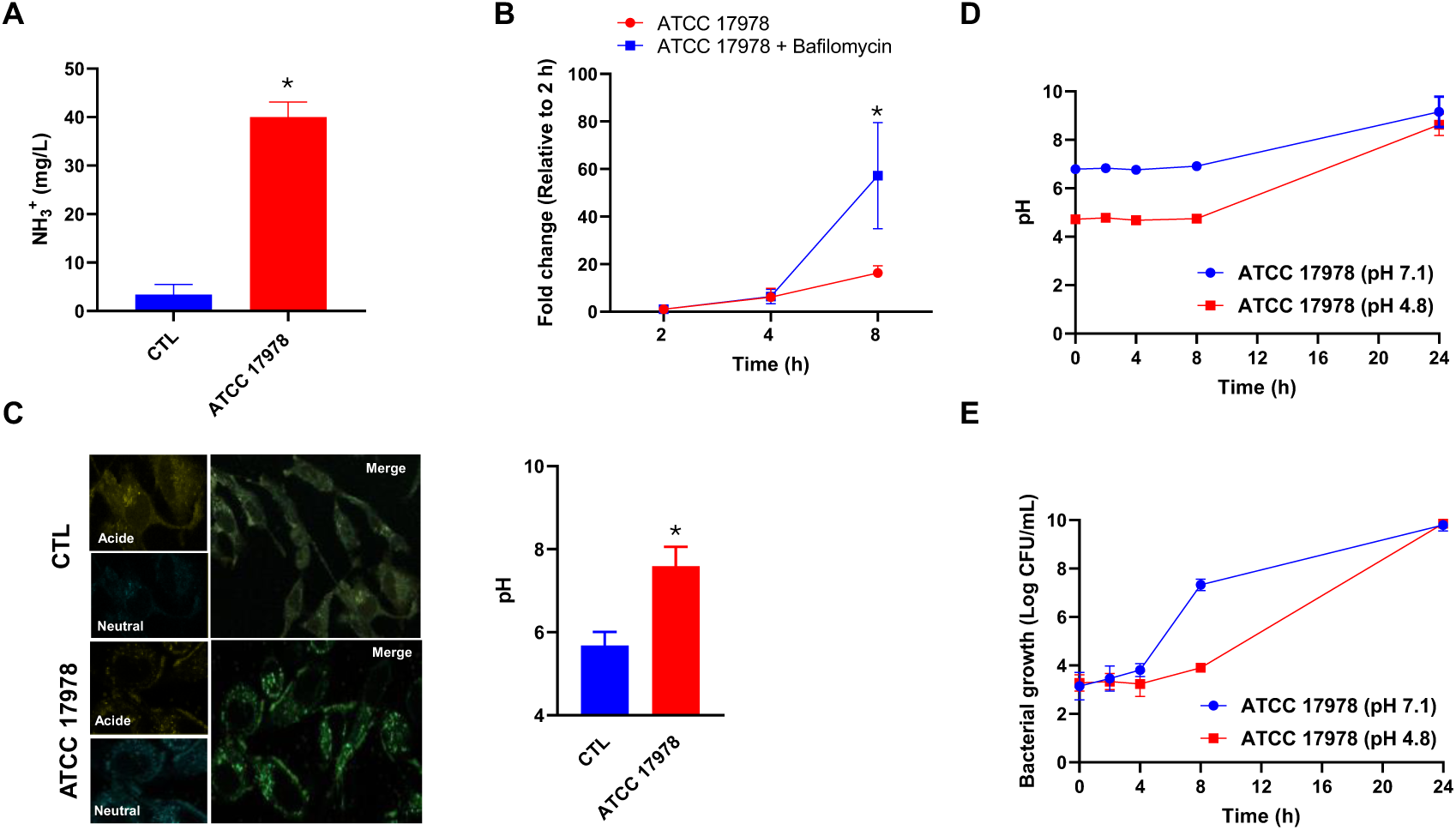
Intracellular survival of *A. baumannii* is supported by ammonia production and pH modulation. **(A)** Ammonia production in HeLa cells infected with the ATCC 17978 strain for 2 h. \**P*=0.011 for ATCC 17978 vs. CTL. **(B)** Intracellular replication of the ATCC 17978 strain in HeLa cells infected at 2, 4 and 8 h post-infection, in the presence or absence of bafilomycin. Data are presented as fold change relative to the intracellular CFU at 2 h. \**P*=0.033 for ATCC 17978 vs. ATCC 17978 + bafilomycin. **(C)** Lysosomal pH measurement of HeLa cells infected for 2 h with the ATCC 17978 strain stained with LysoSensor Yellow/Blue. \**P*=0.0164 for ATCC 17978 vs. CTL. **(D)** pH measurement of LB medium over 24 h in the presence of the ATCC 17978 strain. **(E)** *A. baumannii* ATCC 17978 in LB medium during 24 h under acidic or neutral conditions. CTL: control.

### *In vivo* validation of the iPLA_2_-TFEB axis during *A. baumannii* infection in *C. elegans*

Using *C. elegans* as an *in vivo* model, we monitored activation of HLH-30, the ortholog of human TFEB, in response to *A. baumannii* infection. At 2 and 4 h post-infection, very weak HLH-30 nuclear translocation represented by discrete green fluorescent spots are present, and by 8 and 24 h, there was a high increase in both the intensity and number of fluorescence spot distributed throughout the worm’s body. In contrast, worms fed *E. coli* for 24 h showed significantly less HLH-30 activation (**Figure 6A and B**). These results indicate that *A. baumannii* infection induces a progressive and systemic activation of HLH-30 in *C. elegans*, supporting the involvement of TFEB-dependent transcriptional response during host defense *in vivo*.

**Figure 6.**
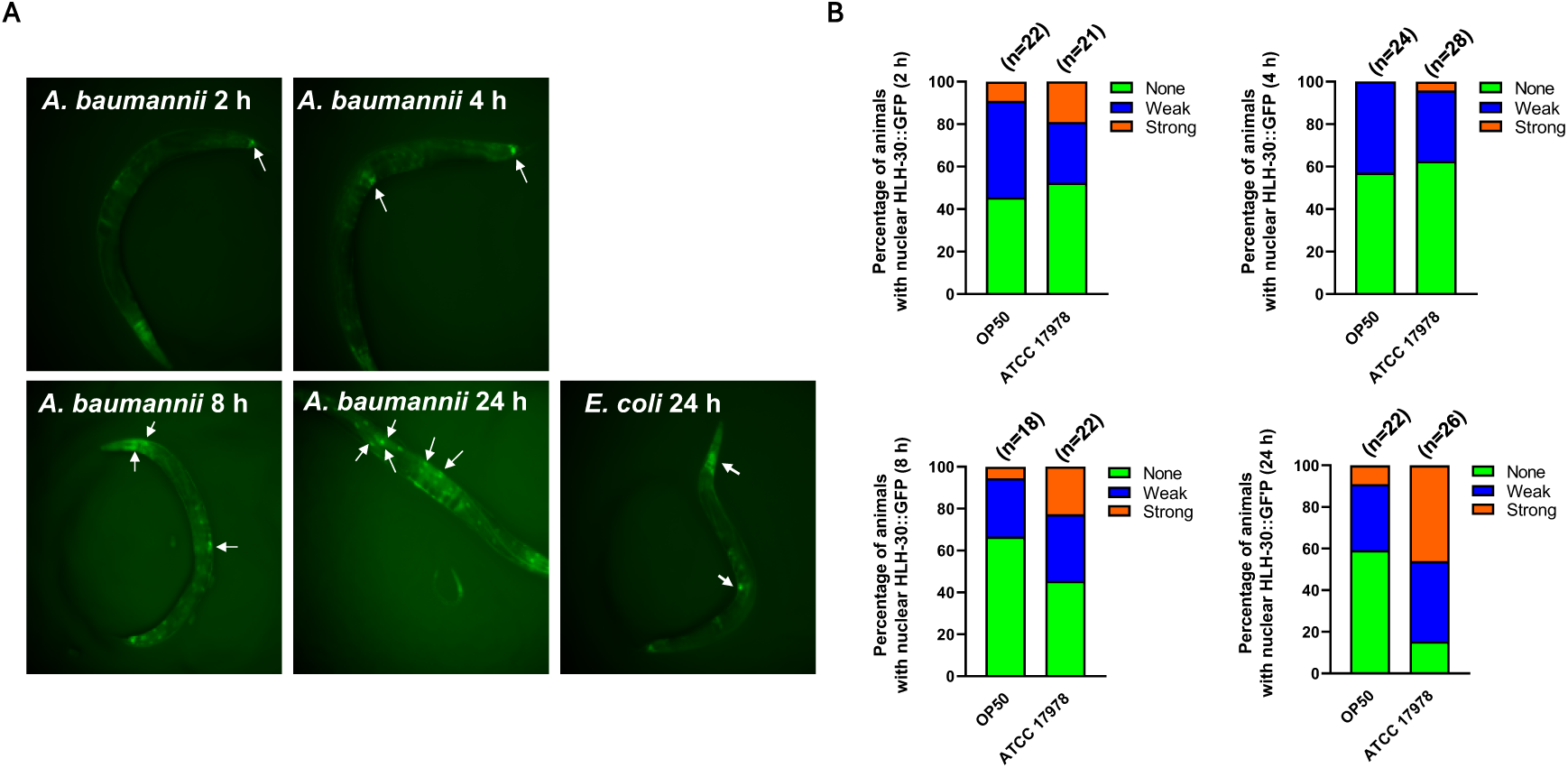
*A. baumannii* increases HLH-30 nuclear translocation in *C. elegans*. (**A**) Fluorescence micrographs and (**B**) quantification of HLH-30::GFP nuclear translocation in tissue cells of transgenic worms expressing the integrated array *sqIs17 [Phlh-30::hlh-30::GFP; rol-6(su1006)]* when grown in presence of the nonpathogenic *E. coli* OP50 or *A. baumannii* ATCC 17978 during 2, 4, 8 and 24 h. n = number of animals.

### OmpA-dependent invasion is required for activation of the iPLA_2_-TFEB signaling pathway

OmpA is a key factor required for successful entry of *A. baumannii* into host cells [Vila Farrès et al. Sci Rep 2017]. To explore whether OmpA-mediated invasion contributes to activation of the iPLA_2_-TFEB signaling axis, we compared the effects of ΔOmpA mutant strain with those of the wild-type strain in HeLa cells. First, we assessed bacterial adherence and invasion efficiency. The ΔOmpA mutant strain exhibited a significant ≥40% reduction in its ability to adhere and invade HeLa cells compared with the wild-type strain (*P*=0.003 and *P*=0.032, respectively) (**Figure 7A**), confirming the importance of OmpA in host cell entry. We next examined the iPLA_2_ enzymatic activity and LPC production. Although the ΔOmpA mutant strain still induced detectable iPLA_2_ enzymatic activity and LPC production, both responses were reduced compared with those induced by the wild-type strain (**Figure 7B and C**). These findings suggest that OmpA-mediated invasion enhances activation of the iPLA_2_ signaling cascade. Finally, we determined TFEB expression following infection with either strain. Total TFEB protein levels were significantly lower in cells infected with the ΔOmpA mutant compared with wild-type infected cells (**Figure 7D**).

**Figure 7.**
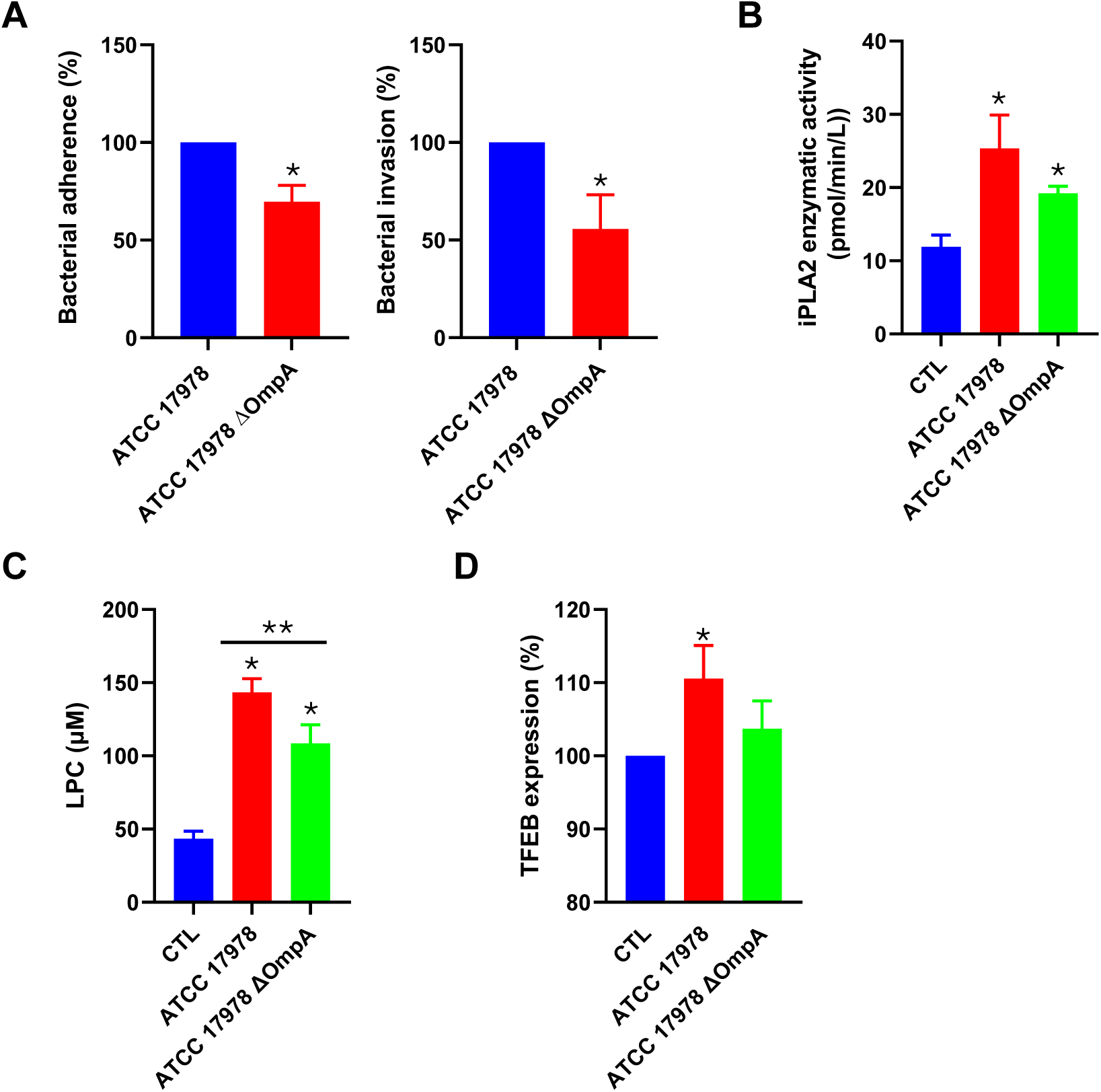
Activation of the iPLA_2_-TFEB pathway depends on OmpA-driven bacterial invasion. **(A)** Adherence and invasion of ATCC 17978 and ATCC 17978 ΔOmpA strains in HeLa cells for 2 h. \**P*=0.003 for ATCC 17978 vs. ATCC 17978 ΔOmpA (adherence), \**P*=0.032 for ATCC 17978 vs. ATCC 17978 ΔOmpA (invasion). **(B)** Enzymatic activity of iPLA_2_ in HeLa cells infected with ATCC 17978 or ATCC 17978 ΔOmpA strains for 24 h. \**P*=0.025 for CTL vs. ATCC 17978, \**P*=0.0255 for CTL vs. ATCC 17978 ΔOmpA. **(C)** LPC production in HeLa cells infected with ATCC 17978 or ATCC 17978 ΔOmpA strains for 24 h. \**P*<0.001 for CTL vs. ATCC 17978, \**P*=0.043 for CTL vs. ATCC 17978 ΔOmpA, \*\**P*=0.03 for ATCC 17978 vs. ATCC 17978 ΔOmpA. **(D)** Immunoblot analysis of TFEB in HeLa cells infected with ATCC 17978 or ATCC 17978 ΔOmpA strains for 2 h. \**P*=0.0424 for CTL vs. ATCC 17978. CTL: control.

In addition, Zhang et al. reported that the metabolite itaconate alkylates TFEB that reduces its cytosolic retention and promotes TFEB-dependent lysosomal biogenesis [Zhang et al. 2022]. Given the structural similarity between itaconate and fumarate [O’Neill and Artymov 2019], we suggest that fumarate might also target TFEB. To test this, we measured fumarate levels in HeLa cells following infection with the ATCC 17978 wild-type strain or the ΔOmpA mutant. Infection with the ATCC 17978 wild type strain increased fumarate levels from 279.53 to 499.50 µg/mL (*P*=0.0423), whereas the ΔOmpA mutant induced a smaller, non-significant increase to 330.32 µg/mL (Figure S3). Collectively, these results indicate that the activation of the iPLA_2_-TFEB pathway is partially dependent on OmpA expression and efficient bacterial invasion, establishing a mechanistic link between *A. baumannii* entry into host cells and induction of host stress and lysosomal defense signaling.

### OmpA is required for lysosomal biogenesis induction and ammonia-mediated intracellular survival of *A. baumannii*

Having demonstrated that *A. baumannii* senses and responds to acidic stress, we sought to determine whether OmpA is required for the induction of lysosomal biogenesis, ammonia production and neutral pH during infection. Fluorescence microscopy analysis revealed that infection with the ΔOmpA mutant strain failed to induce significant lysosomal labelling. Lysosomal labeling returned to near-baseline levels (approximately 20%), comparable to uninfected control cell, whereas the wild-type strain induced robust lysosomal labeling (approximately 70%; *P*<0.014) (**Figure 8A and B**). These findings indicate that OmpA is necessary for efficient activation of the host lysosomal response. We next measured ammonia production in HeLa cells and in bacterial culture medium and found that the ΔOmpA mutant produced significantly lower levels of ammonia (15 mg/L) compared with the wild-type strain (40 mg/L; *P*=0.033) in HeLa cells (**Figure 8C**) and in bacterial culture medium (98.79 mg/L *vs*. 49.85 mg/L; *P*=0.0389) (**Figure S4**), suggesting that OmpA contributes to bacterial ammonia generation under intracellular conditions. Consistently, infection with the ΔOmpA mutant resulted in a lower lysosomal (pH from 7.6 to 6.7) (**Figure 8D**). Furthermore, when host lysosomal acidification was inhibited using bafilomycin, replication of the ΔOmpA mutant strain was markedly impaired compared with the wild-type strain (**Figure 8E**). Collectively, these results demonstrate that OmpA coordinates host-*A. baumannii* interactions by promoting bacterial invasion, activating iPLA_2_-TFEB axis, inducing lysosomal biogenesis, and supporting ammonia-mediated adaptation to acidic stress.

**Figure 8.**
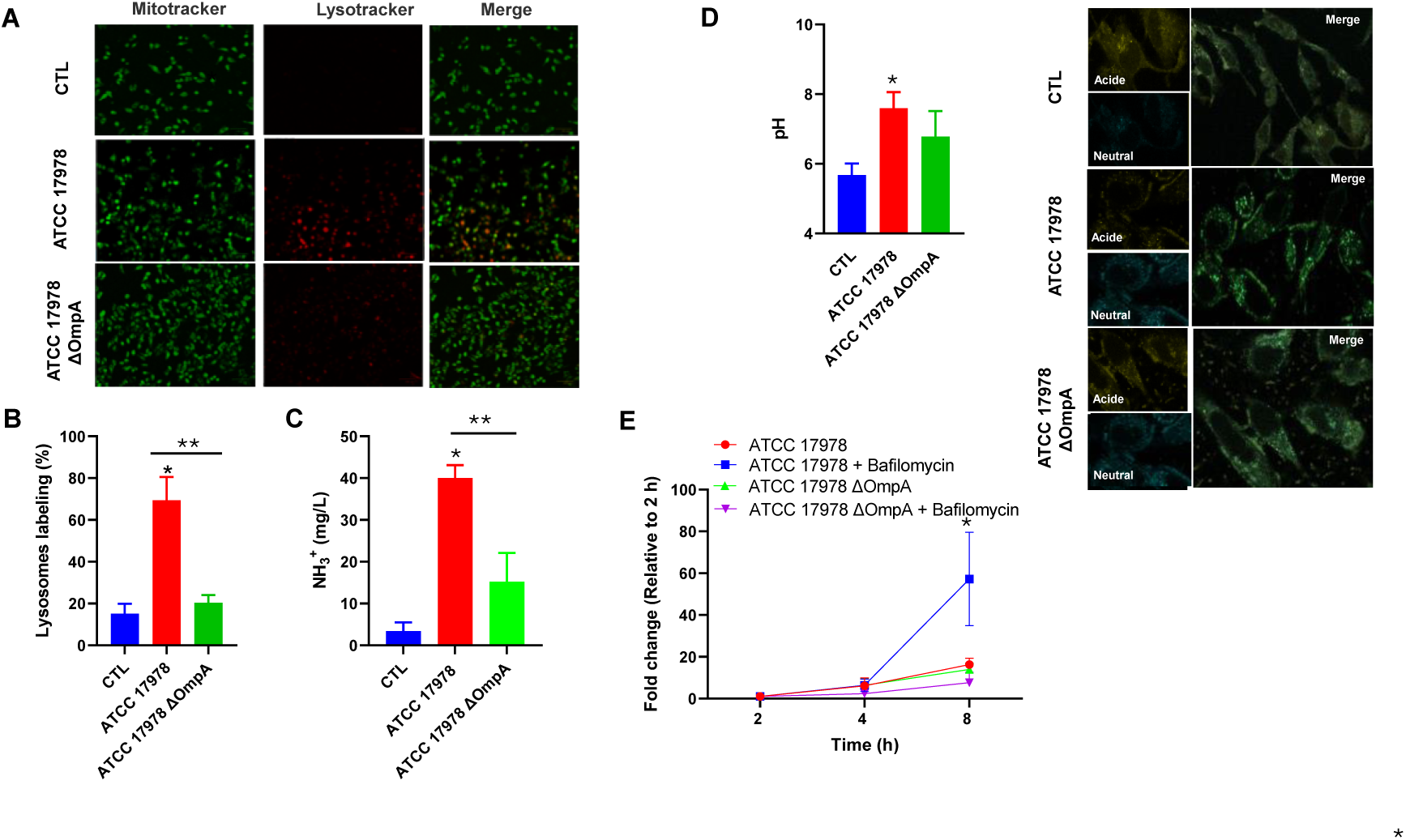
OmpA mediates lysosomal biogenesis induction and ammonia-dependent intracellular survival of *A. baumannii*. **(A and B)** Lysosomal analysis in HeLa cells infected with ATCC 17978 or ATCC 17978 ΔOmpA strains for 2 h. Acidic organelles were labeled with LysoTracker Red and mitochondria were labeled with MitoTracker Green. Lysosomal staining in infected HeLa cells is presented in the bar graph as a percentage relative to non-infected control cells. \**P*=0.0111 for ATCC 17978 vs CTL. \*\**P*=0.0140 for ATCC 17978 vs. ATCC 17978 ΔOmpA. **(C)** Ammonia production in HeLa cells infected with ATCC 17978 or ATCC 17978 ΔOmpA strains for 2 h.\**P*=0.011 for ATCC 17978 vs CTL, \*\**P*=0.03031 for ATCC 17978 vs ATCC 17978 ΔOmpA. **(D)** Intracellular replication of ATCC 17978 or ATCC 17978 ΔOmpA strains in HeLa cells infected at 2, 4 and 8 h post-infection, in the presence or absence of bafilomycin. Data are expressed as fold change in intracellular CFU relative to 2 h.\**P*=0.033 for ATCC 17978 vs. ATCC 17978 + bafilomycin. **(E)** Lysosomal pH measurement of HeLa cells infected with the ATCC 17978 or ATCC 17978 ΔOmpA strains stained with LysoSensor Yellow/Blue dextran for 2 h. \**P*=0.0164 for ATCC 17978 vs. CTL. CTL: control.

### OmpA contributes to HLH-30 nuclear translocation in *C. elegans*

To address the role of OmpA in HLH-30 nuclear translocation in the context of an infective process in a complete organism, we assessed HLH-30 activation in *C. elegans* exposed to *A. baumannii*. We suggest that OmpA is required for *A. baumannii* to induce HLH-30 nuclear translocation. To test this, we determined HLH-30 nuclear translocation in worms infected with ATCC 17978 wild-type and ΔOmpA mutant strain. Interestingly, the ΔOmpA mutant induced significantly very weak and less strong HLH-30 nuclear transclocation compared with the wild-type strain (**Figure 9**).

**Figure 9.**
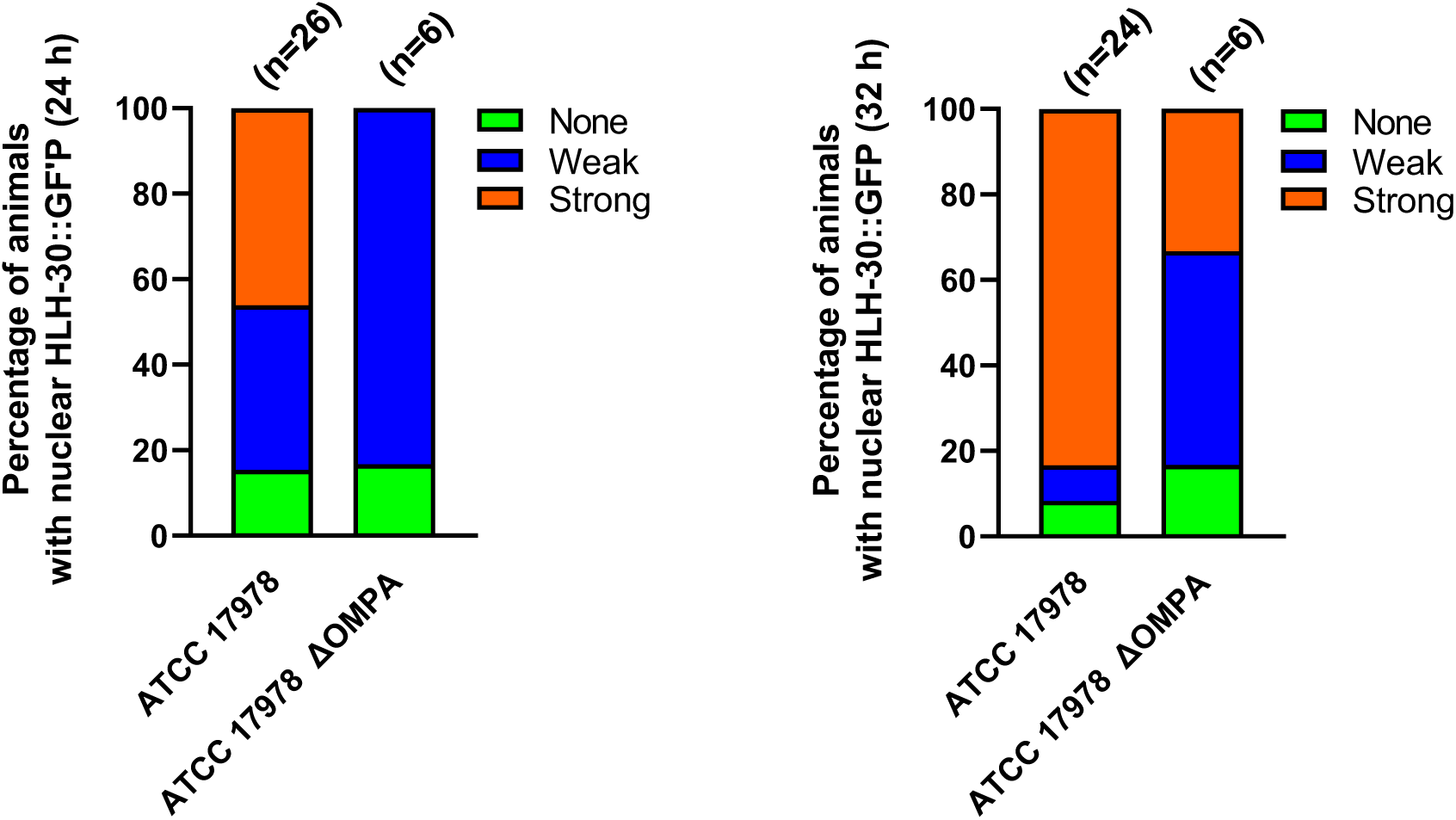
OmpA is involved in HLH-30 nuclear translocation in *C. elegans*. Quantification of HLH-30::GFP nuclear translocation in tissue cells of transgenic worms expressing the integrated array *sqIs17 [Phlh-30::hlh-30::GFP; rol-6(su1006)]* when grown in presence of *A. baumannii* ATCC 17978 or ATCC 17978 ΔOmpA during 24 and 32 h. n = number of animals.

## Discussion

In this study, we identify a previously unrecognized signaling axis linking *A. baumannii* infection to host lipid signaling, calcium influx, and lysosomal regulation. Our data demonstrate that *A. baumannii* activates iPLA_2_, leading to increased Cox-2 activity and PGE2 production, and that this lipid signaling cascade functions upstream of TFEB activation. These finding expand the current understanding of host-pathogen interactions by positioning iPLA_2_ as a central mediator that connects bacterial invasion to lysosomal adaptive responses. A mechanism we previously suggested in earlier studies [Smani et al JBC 2012, Parra Millán 2018].

The observed increase in iPLA_2_ expression and enzymatic activity following infection indicates that *A. baumannii* triggers a host stress response associated with membrane phospholipid remodeling. The attenuation of Cox-2 and PGE2 production by the irreversible iPLA_2_ inhibitor BEL supports that activation of this inflammatory branch is largely iPLA_2_-dependent. For instance, iPLA_2_ activation by effector caspases has been reported to hydrolyse membrane bound phosphatidylcholine to bioactive LPC and arachidonic acid [Lauber et al. Cell 2003]. This last is converted by Cox-2 to produce PGE2 [Flower Nat Rev Drug Discov 2023]. Given the well-established roles of PGE2 in modulating inflammation and immune signaling, activation of the iPLA_2_-Cox-2-PGE2 may contribute both to host defense and to *A. baumannii*-driven immune modulation, as suggested by our previous studies [Smani et al AAC 2015, Miró Canturri et al 2021]. Importantly, we demonstrate that iPLA_2_ regulates TFEB expression and nuclear translocation. Pharmacological inhibition and siRNA-mediated knockdown consistently reduced TFEB levels, supporting a direct regulatory link. Mechanistically, our results indicate that LPC, generated downstream of iPLA_2_ activation [Smani et al Nat Bio Cell 2024], acts as a bioactive lipid mediator promoting TFEB activation. This finding integrates lipid metabolism with lysosomal transcription control and suggests that membrane-derived lipid signals serve as upstream modulators of autophagic and lysosomal programs during bacterial infection. Phagosome maturation by LPC has been reported in macrophages infected by *Mycobacterium tuberculosis* [Lee et al. 2018] We further show that calcium influx through ORAI1 is required for TFEB activation, consistent with the established role of calcium-calcineurin signaling in TFEB nuclear translocation. Medina *et al*. revealed that lysosomes can release calcium via MCOLN1 under starvation conditions. This lysosomal calcium efflux activates the phosphatase calcineurin, which desphosphorylates TFEB, thereby promoting its nuclear translocation and autophagy [Medina et al 2015]. In the present study, both pharmacological inhibition and genetic silencing of ORAI1 significantly impaired TFEB activation and lysosomal biogenesis, establishing ORAI1-mediated calcium entry as a critical component of this pathway. However, Sbano et al (2017) reported that TFEB overexpression in HeLa cells reduces calcium influx through SOCE and decreases calcium refilling in the endoplasmic reticulum of HeLa cells. Based on these observations, we propose that enhanced TFEB activation following ORAI1-dependent calcium influx may trigger a negative feedback loop that modulates the TFEB-mediated regulation of lysosomal biogenesis. Collectively, our findings position calcium signaling downstream of iPLA_2_/LPC and upstream of TFEB, suggesting the existence of a coordinated lipid-calcium signaling axis controlling lysosomal function and autophagy.

Despite enhanced lysosomal biogenesis, *A. baumannii* efficiently survives intracellularly. Our findings suggest that ammonia production in host cells and environmental alkalinization represent key bacterial countermeasures against host lysosomal acidification. *A. baumannii* is able to tolerate acidic conditions and continue to grow, as we previously reported [Parra Millán et al. 2018] by alkalinizing the culture medium through ammonia production. Some studies have reported that Gram-negative bacteria produce ammonia within host cells, particularly in macrophages [Distel et al 2023, Huang et al. 2022]. However, one report described that the *A. baumannii* ABC141 strain does not secrete ammonia in endothelial cells [Debruyne et al 2026]. The marked increase in bacterial replication following treatment with bafilomycin further underscores the importance of lysosomal acidification in restricting bacterial growth. The ability of *A. baumannii* to actively increase extracellular pH highlights its metabolic adaptability and its capacity to neutralize acidic stress.

The *in vivo* validation in *C. elegans* strengthens the physiological relevance of the iPLA_2_-TFEB axis. Activation of the HLH-30, the TFEB ortholog, during *A. baumannii* infection supports the evolutionary conservation of this lysosomal transcriptional response and suggests that this pathway represents a fundamental host defense mechanism. Other evolutionarily conserved pathways involved in host defense against bacterial infection have also been described, such as the phospholipase C-protein kinase D-TFEB signaling axis [Najibi et al 2016].

Our data also identify OmpA-dependent invasion as a critical upstream event required for full activation of the iPLA_2_-TFEB pathway. The ΔOmpA mutant exhibited reduced cellular invasion, diminished iPLA_2_ activity, LPC and fumarate production, impaired TFEB activation, decreased lysosomal biogenesis, and lower ammonia production. OmpA is widely recognized as an important virulence factor in several microorganisms, including *A. baumannii*, where it has been shown to activate effector caspases, precursor of iPLA_2_, both *in vitro* and *in vivo* [Choi et al. 2005, Lee et al 2010, Zhao et al. 2026]. Importantly, one study reported that OmpA from *A. baumannii* promotes autophagy in rat lung epithelial cells through modulation of the mTOR pathway *in vivo* [Zhao et al 2021]. Notably, mTOR-dependent phosphorylation is a key regulator of TFEB nuclear translocation [Napolitano et al 2018]. In addition, we showed that the ΔOmpA mutant induced lower fumarate production. Fumarate, a tricarboxylic acid metabolite, has been reported to enhance phagocytosis in macrophages of fish [Yang et al Font Mol Biosci 2021]. Together, these observations suggest that OmpA may influence host defense responses not only by modulating autophagy through the mTOR-TFEB axis but also by affecting metabolite-mediated enhancement of phagocytosis.

Moreover, OmpA from other microorganism, such as *M. tuberculosis*, has been shown to accelerate ammonia secretion and facilitate bacterial adaptation to acidic environments [Song et al. 2011]. Ammonia production can inhibit phagosome-lysosome fusion in macrophages [Gordon et al 1980], a process that has also been reported to be disrupted *A. baumannii* OmpA [An et al 2019]. Collectively, these findings establish a mechanistic link between bacterial entry and host lipid-lysosomal signaling responses. OmpA therefore appears to coordinate both the activation of host signaling pathways and bacterial adaptive survival strategies that promote intracellular survival.

In summary, this study defines a novel OmpA-iPLA_2_-LPC-ORAI1-TFEB signaling axis activated during *A. baumannii* infection. While this pathway promotes lysosomal biogenesis as a host defense response, *A. baumannii* simultaneously deploys ammonia-mediated alkalinization to counteract lysosomal killing. These findings provide new insight into the dynamic interplay between host stress signalling and bacterial adaptation and may open avenues for therapeutic strategies targeting iPLA_2_-TFEB axis to enhance intracellular bacterial clearance.

## ACKNOWLEDGMENTS

This research was funded by the Ministerio de Ciencia e Innovación, Agencia Estatal de Investigación, Fondo Europeo de Desarrollo Regional, MCIN/AEI/10.13039/501100011033/FEDER, UE (Grant PID2022-136357OBI00), by the Consejería de Universidad, Investigación e Innovación de la Junta de Andalucía (Grant ProyExcel_00116), and by VI Plan Propio de Investigación y Transferencia 2023-2026 de la Universidad Pablo de Olavide (Ayuda A3, Referencia: PP12303). C.A.R is supported by V Plan Propio de Investigación y Transferencia 2018-2020 de la Universidad Pablo de Olavide (Ayuda B2, Referencia: PP12205).

We thank Giulia Olgiati, Moisés Pérez Pérez and Laura Tomás for their technical help and Tanja Dapa and Tarik Smani for reviewing the manuscript.

## AUTHOR CONTRIBUTIONS

**Irene Molina Panadero**: Methodology; Investigation, Formal analysis. **Angela Rey Hidalgo**: Methodology; Investigation; Formal analysis. **María José López Carballo**: Methodology; Investigation; Formal analysis. **Celia Atalaya Rey**: Methodology; Investigation, Formal analysis. **Manuel J. Muñoz Ruiz**: Investigation; Writing-Review and Editing. **Younes Smani**: Writing-Review and Editing; Supervision; Funding Acquisition; Conceptualization.

## DATA AVAILABILITY

The data supporting the findings of this study are available from the corresponding author upon reasonable request.

---

**Figure S1.**
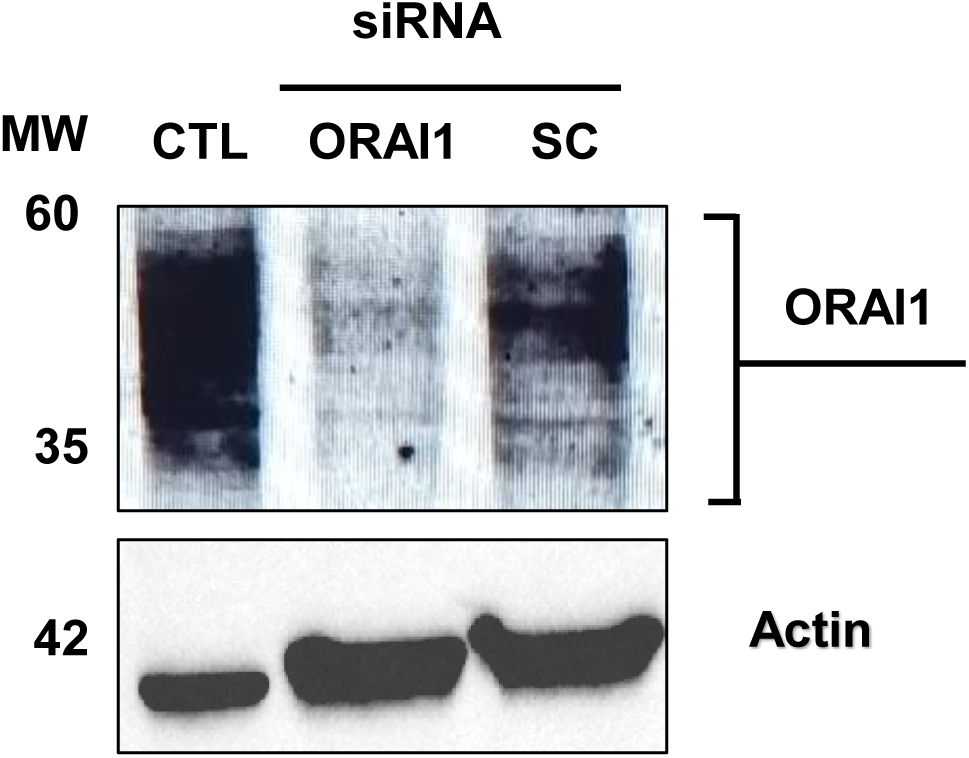
Immunoblot analysis of HeLa cells transfected with scrambled (SC) and ORAI siRNA for 48 h.

**Figure S2.**
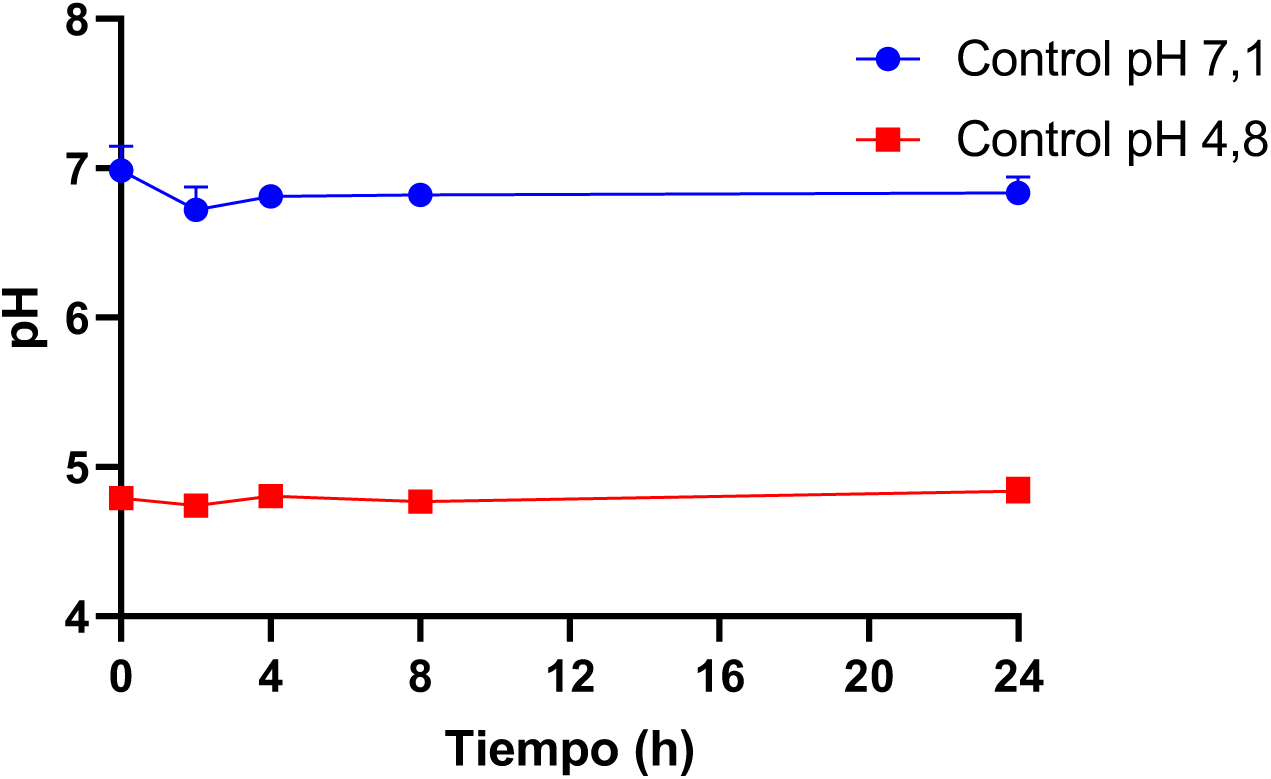
pH determination during 24 h of LB medium without *A. baumannii* ATCC 17978 strain.

**Figure S3.**
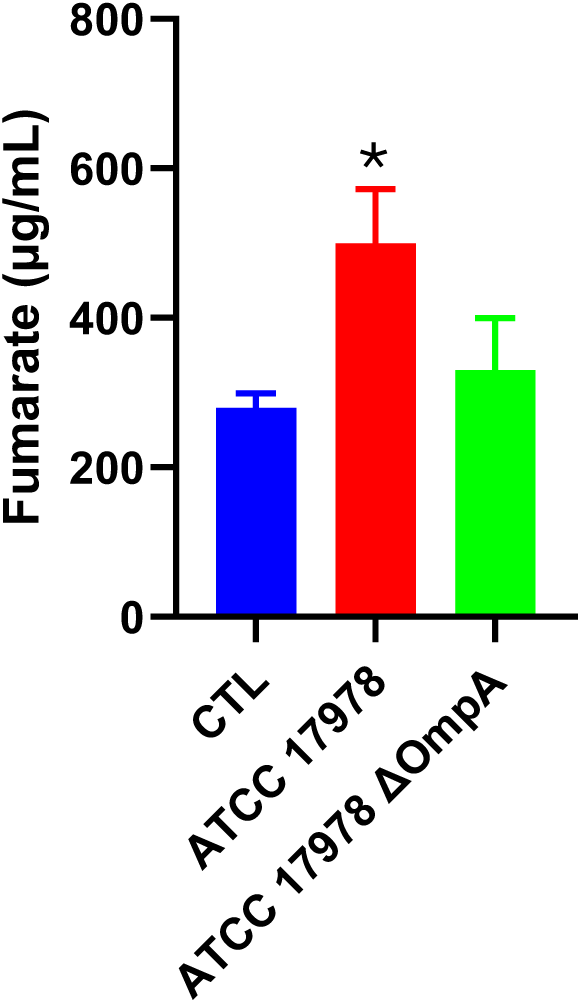
Fumarate production in HeLa cells infected with ATCC 17978 or ATCC 17978 ΔOmpA strains for 2 h.\**P*=0.0423 for ATCC 17978 vs CTL. CTL: control.

**Figure S4.**
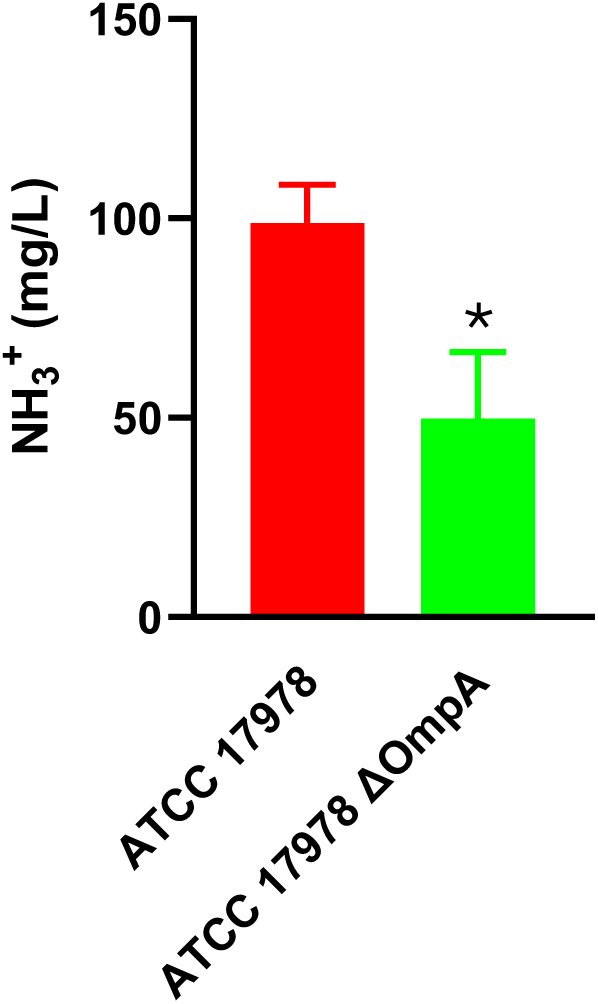
Ammonia production by *A. baumannii* ATCC 17978 and ΔOmpA mutant strains in bacterial medium during 2 h of growth. \**P*=0.0389 ATCC 17978 vs **Δ**OmpA.

